# Online biophysical predictions for SARS-CoV-2 proteins

**DOI:** 10.1101/2020.12.04.411744

**Authors:** Luciano Kagami, Joel Roca-Martínez, Jose Gavaldá-García, Pathmanaban Ramasamy, K. Anton Feenstra, Wim Vranken

**Affiliations:** Interuniversity Institute of Bioinformatics in Brussels, ULB-VUB, Triomflaan, Brussels 1050, Belgium; Structural Biology Brussels, Vrije Universiteit Brussel, Pleinlaan 2, Brussels 1050, Belgium; VIB Structural Biology Research Centre, Pleinlaan 2, Brussels 1050, Belgium; VIB-UGent Center for Medical Biotechnology, VIB, Ghent 9000, Belgium; Department of Biomolecular Medicine, Faculty of Health Sciences and Medicine, Ghent University, Ghent 9000, Belgium; IBIVU – Center for Integrative Bioinformatics, Vrije Universiteit Amsterdam, Amsterdam 1081HV, The Netherlands; AIMMS – Amsterdam Institute for Molecules Medicines and Systems, Vrije Universiteit Amsterdam, Amsterdam 1081HV, The Netherlands

**Keywords:** Proteins, single sequence based predictions, biophysical features, SARS-CoV-2, COVID-19

## Abstract

The SARS-CoV-2 virus, the causative agent of COVID-19, consists of an assembly of proteins that determine its infectious and immunological behavior, as well as its response to therapeutics. Major structural biology efforts on these proteins have already provided essential insights into the mode of action of the virus, as well as avenues for structure-based drug design. However, not all of the SARS-CoV-2 proteins, or regions thereof, have a well-defined three-dimensional structure, and as such might exhibit ambiguous, dynamic behaviour that is not evident from static structure representations, nor from molecular dynamics simulations using these structures. We here present a website (http://sars2.bio2byte.be/) that provides protein sequence-based predictions of the backbone and side-chain dynamics and conformational propensities of these proteins, as well as derived early folding, disorder, β-sheet aggregation and protein-protein interaction propensities. These predictions attempt to capture the ‘emergent’ properties of the proteins, so the inherent biophysical propensities encoded in the sequence, rather than context-dependent behaviour such as the final folded state. In addition, we provide an indication of the biophysical variation that is observed in homologous proteins, which give an indication of the limits of the functionally relevant biophysical behaviour of these proteins. With this website, we therefore hope to provide researchers with further clues on the behaviour of SARS-CoV-2 proteins.

## Introduction

The SARS-CoV-2 virus, the causative agent of COVID-19, consists of an assembly of proteins that determine its infectious and immunological behavior, as well as its response to therapeutics. Major structural biology efforts on these proteins have already provided essential insights into the mode of action of the virus, as well as avenues for structure-based drug design. However, not all of the SARS-CoV-2 proteins, or regions thereof, have a well-defined three-dimensional structure, and as such might exhibit ambiguous, dynamic behaviour that is not evident from static structure representations generated by structural biology approaches, nor from molecular dynamics simulations using these structures.

We here present a website (http://sars2.bio2byte.be/) that provides extensive protein sequence-based predictions for the SARS-CoV-2 proteins, which can help to identify behavior or features of these proteins that are might not be captured by structural biology or molecular dynamics approaches. The predictions include the DynaMine backbone^1,2^ and side-chain dynamics^3^ as well as conformational propensities^3^, and derived DisoMine disorder^4^, EFoldMine early folding^3^, Agmata β-sheet aggregation^5^, SeRenDIP protein-protein interaction^6^ and SeRenDIP-CE conformational epitope propensities^7^. These predictions attempt to capture the ‘emergent’ properties of the proteins, so the inherent biophysical propensities encoded in the sequence, rather than context-dependent behaviour such as the final folded state. This approach has already shown promise in, for example, detecting remote homologues by biophysical similarity, which gives more accurate results than directly using amino acid information^8^. We apply this concept on the SARS-CoV-2 proteins by incorporating evolutionary information, so enabling us to display the biophysical variation observed in homologous proteins, which indicates likely limits of the functionally relevant biophysical behaviour of these proteins.

## Methods

### Datasets

The target amino acid sequences of the 14 proteins were obtained from the UniProt^9^ COVID-19 section, after filtering on ‘Other organisms’ by ‘Severe acute respiratory syndrome coronavirus 2’. Multiple sequence alignments (MSAs) for these sequences were obtained using a BLAST search from UniProt using default parameters against the Uniref90 protein dataset. This was followed by the standard UniProt ClustalW alignment procedure to obtain the MSA.

### Predictions

On each target sequence, the backbone dynamics (DynaMine)^1,2^, and related side-chain dynamics and conformational propensities^3^ were predicted at the per-amino acid level, as well as early folding (EFoldMine)^3^, disorder (DisoMine)^4^, β-sheet aggregation (Agmata)^5^, protein-protein interactions (SeRenDIP)^6,10^, and SeRenDIP-CE epitope propensities ^7^. Predictions of FUS-like phase separation were also performed with PSPer^11^. All predictions except for Agmata, SeRenDIP, SeRenDIP-CE and PSPer were then run on all individual sequences in the MSAs, with the values mapped back to the MSA, so obtaining per MSA column a list of prediction values. A standard box plot approach was then applied to each per-column list of values to identify the median, first and third quartile, and outlier range per biophysical feature per column in each MSA.

### Phosphorylation sites

Experimentally validated phosphorylation sites were obtained from two SARS-CoV-2 phosphoproteome projects (PXD020183, PXD019113) in the PRIDE repository^12^. The search files for the projects were downloaded and processed to extract the phospho-site information. Since the data processing protocol varies between the projects depending on the search engine used, we only considered the phosphor-sites that are seen in more than one project with a localization probability of >0.6.

### Website

The information was visualized online using the Django framework, with the ApexCharts JavaScript library employed for visualization of the predictions and their MSA distribution.

## Results

### Website description

The home page provides a brief statement on the purpose of the website, and how to proceed. On the ‘Entries’ page (available from the top bar), each SARS-CoV-2 protein is listed by its UniProt identifier, with its UniProt-based description provided on the right-hand side of the page. For each entry, the following links are provided:

- Sequence-based predictions (click on the UniProt ID)
- Structure(s) in the PDB^13^ via the PDBe-KB^14^ (if available)
- UniProt information
- PSPer predictions about the possible phase-separation behavior of this protein
- Download all predictions for this protein in JSON format

The per-protein pages provide first a visualization of all incorporated predictions (y-axis) in function of the protein sequence (x-axis). Hovering with the pointer over the graph will display the residue number (below the x-axis) and the corresponding prediction values (in the legend). The following sequence-based predictions are provided, in different colors:

- **DynaMine backbone dynamics** (black): values above 0.8 indicate rigid conformations, values above 1.0 membrane spanning regions, values below 0.69 flexible regions. Values between 0.80-0.69 are ‘context’ dependent and capable of being either rigid or flexible.
- **DynaMine sidechain dynamics** (grey): Higher values mean more likely rigid. These values are highly dependent on the amino acid type (*i*.*e*. a Trp will be rigid, an Asp flexible).
- **DynaMine conformational propensities (sheet, helix, coil)** (blue, red, purple): Higher values indicate higher propensities.
- **EFoldMine earlyFolding propensity** (green): Values above 0.169 indicate residues that are likely to start the protein folding process, based on only local interactions with other amino acids.
- **DisoMine disorder** (yellow): values above 0.5 indicate that this is likely a disordered residue.
- **Agmata aggregation propensity** (dark green): These values are divided by a factor of 20 from the original. Peaks indicate residues likely to be involved in β-sheet aggregation.
- **SeRenDIP protein-protein interactions (PPI)** (cyan): values above 0.5 indicate that this residue likely participates in protein-protein interactions.
- **SeRenDIP-CE conformational epitope regions (CE)** (seagreen): values above 0.5 indicate that this residue likely is part of a conformational (discontinuous) epitope region.

These predictions reflect ‘emerging’ properties, so what the sequence is capable of, not necessarily what it will adopt in a final fold. Each prediction can be toggled on/off by clicking on the corresponding name in the legend of the plot.

The second plot provides a visualization of the MSA-based variation of a specific predicted feature (like backbone dynamics) for single-sequence based predictions, again with the prediction value (y-axis) in function of the protein sequence (x-axis). The type of prediction shown can be selected using the ‘Select prediction’ selection box, with the plot showing median (black), first and third quartile (dark grey) and outlier range (light grey) of the distribution per column in the MSA, as well as the original prediction for the target protein itself (red), which corresponds to the prediction in the top graph. Note that only MSA columns for which there is no gap in the target sequence are shown. These distributions reflect the ‘evolutionary allowed’ range of the biophysical features, which as we have previously shown tends to be only weakly correlated with amino acid sequence-based MSA measures such as entropy^6,15^. Values of the red line outside of the quartile range therefore indicate rather unusual behaviour for this particular protein compared to its homologues, and might indicate interesting areas where this SARS-CoV-2 protein differs from other proteins.

Finally, at the bottom of the page links are provided to the PSPer predictions and the JSON containing all the prediction values, as well as distributions, for this protein.

### Use-case example

The P0DTC9 protein is a nucleoprotein of 419 amino acids with both monomeric and oligomeric forms that interact with RNA, as well as with protein M and NSP3. These interactions tether the genome to the newly translated replicase-transcriptase complex at a very early stage of infection. Structural information is available for UniProt-numbered residues Gly44-Ser180 (Figure 1, box A, based on PDB codes 6yi3, 7act, 7acs), which mediates RNA binding, and for Thr247-Pro364 (Figure 1, box B, based on PDB code 6zco), which are involved in oligomerisation (see also https://www.ebi.ac.uk/pdbe/pdbe-kb/covid19/P0DTC9). The predictions for this protein, displayed in Figure 1 (for an interactive version, please see http://bio2byte.be/sars2/prediction/P0DTC9), show a high propensity for disorder throughout the protein, with backbone dynamics also indicating overall high flexibility (values below 0.69), except for the previously mentioned Gly44-Ser180 and Thr247-Pro364 regions, which have been observed to fold. The N-terminal region prior to Gly44 is highly flexible, with some helix and sheet propensity, and a propensity for protein-protein interactions, but with no indications of early folding or aggregation. It also contains multiple confirmed phosphorylation sites (Ser23 and Ser26), hinting at a possible regulation role.

**Figure 1:**
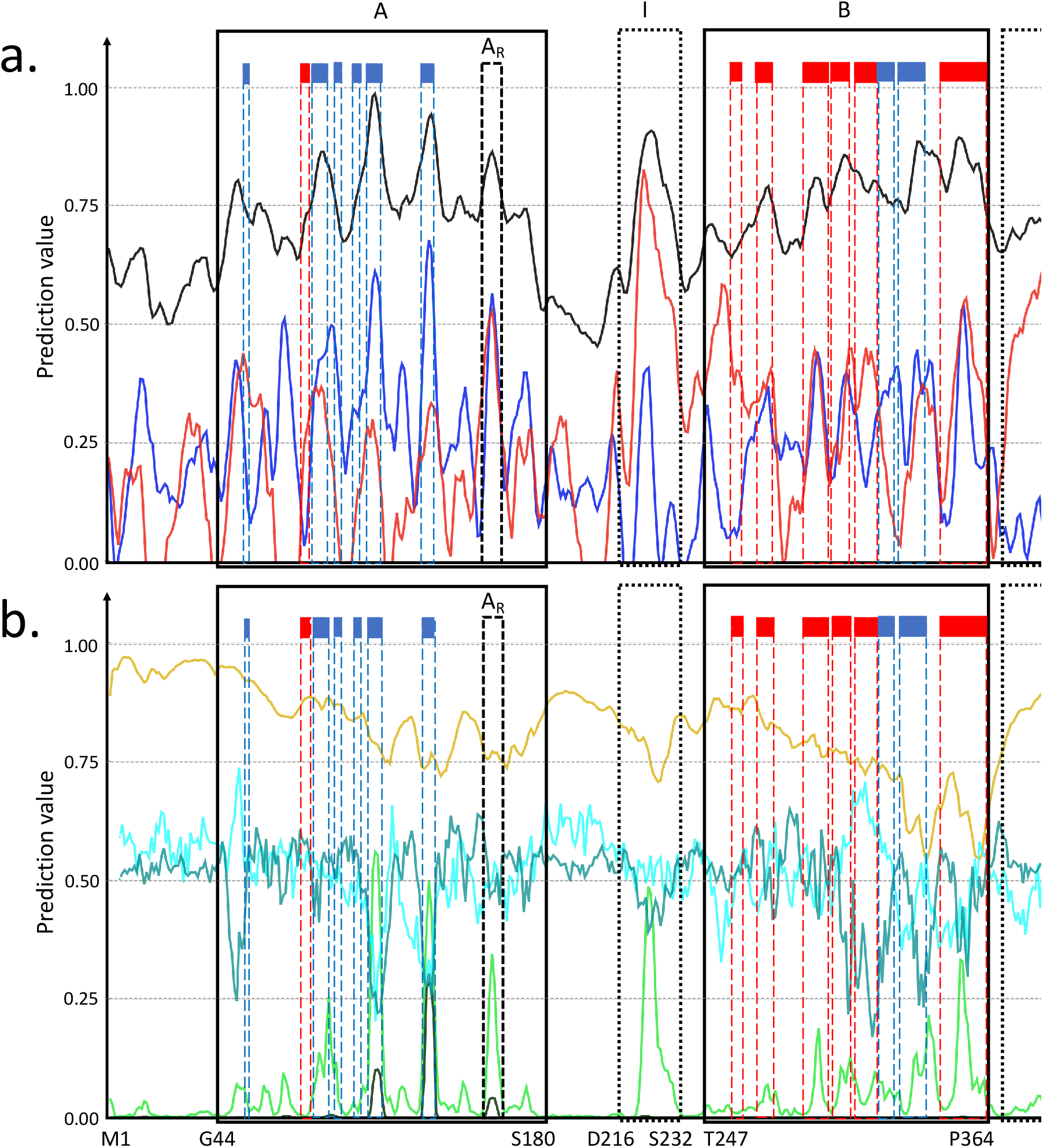
Predictions for the P0DTC9 SARS-CoV-2 protein amino acids (x-axis) for a) backbone dynamics (black), helix (red) and sheet (blue) propensity, and b) early folding (light green), disorder (yellow) β-sheet aggregation (dark green), protein interaction (cyan) and epitopes (seagreen). The two regions for which structures have been determined are indicated by black boxes (A, B), with annotations for consensus α-helix (red boxes) and β-strands (blue boxes) based on these structures included. Regional highlights not evident in these structures (A_R_, I, II) are discussed in the text. On the interactive plots on the server, predictions can be toggled on and off by clicking on their name.

For the first folded domain (box A), the regions confirmed by PBDe-KB to form α-helices (red dotted boxes) and β-strands (blue dotted boxes) are indicated, which tend to correspond to rigid areas with strong secondary structure propensity. Interestingly, the A_R_ region from Asn153 to Gln 163 (black box) does not have a regular secondary structure, but corresponds to an extended region that loops over the outside of the protein. Given the very high prediction values for rigidity, helix and sheet propensity (equal) and early folding, combined with a peak in aggregation, this region could have an important role in the folding process and overall behavior of this protein, even though it does not particularly stand out in the solved crystal structures. There is also notable aggregation tendency corresponding to the 5^th^ and 6^th^ β-strands, and a high epitope propensity in the subsequent region between 6^th^ β-strands and the A_R_ region. A confirmed phosphorylation site is Ser79, at the beginning of the first α-helix.

The subsequent region between Ser180 and Thr247 contains a region with relatively consistent properties from Ser180-Gly215, indicating a highly flexible linker that connects the two structured regions, but with an interestingly elevated PPI propensity. This area also contains multiple confirmed phosphorylation sites (Ser187, Ser194, Ser197, Ser201, Ser206), indicating a regulatory role. The region from Asp216-Thr247 (box I), on the other hand, shows strong peaks in both backbone rigidity and helical propensity from Asp216-Ser232, with indications that this region is prone to early folding. To the best of our knowledge, no structural or functional information is available for this region, but the predictions again indicate that this area could well play a role in regulation, for example, by blocking a site when this helix is formed, or by constraining the distance between the two domains by adapting the overall linker length. Noteworthy is also that both the backbone dynamics and helical propensity, but especially the early folding, are above the third quartile range observed in homologous proteins (Figure S1, S2), indicating that this region has a stronger tendency to autonomously form a helix in the SARS-CoV-2 protein compared to its close homologues.

The oligomerisation domain (box B) shows a strong epitope propensity from Ala273-Asn285, and a very strong PPI propensity from Lys299-Met322, corresponding to the 6^th^ α-helix and 7^th^ β-strand, in line with orientation in the dimer where β-strands 7 and 8 form a four-stranded sheet with the corresponding strands from the other monomer, with α-helix 6 below the sheet and also part of the homodimer interface ^16^. The conformational preference for helix formation is already indicated by the predictions, as are the two β-strand regions.

Finally, the C-terminal region after Pro364 (box II) is in the ‘context-dependent’ zone of the backbone dynamics predictions between 0.80 and 0.69, indicating it could fold, in this case likely into a helix as it also has a strong helical propensity. This again indicates a possible regulatory or transient binding role, possibly to a protein as it has peaks of fairly high PPI propensity. There are also high peaks of epitope propensity in this region, particularly around Pro365 and Leu395-Gln408. The region also contains multiple likely phosphorylation sites.

## Discussion

With this website, approved as an ELIXIR-Belgium emerging service in 2020, we provide researchers with information on possible behaviours of SARS-CoV-2 proteins that are not evident from the static models generated by structural biology, nor from molecular dynamics simulations based on these models. It enables the exploration of these proteins from a different perspective and should help our further understanding of the mode of action of the overall virus.

