## Supplementary material for "Online biophysical predictions for SARS-CoV-2 proteins": Figure S1, S2

### Supplementary data

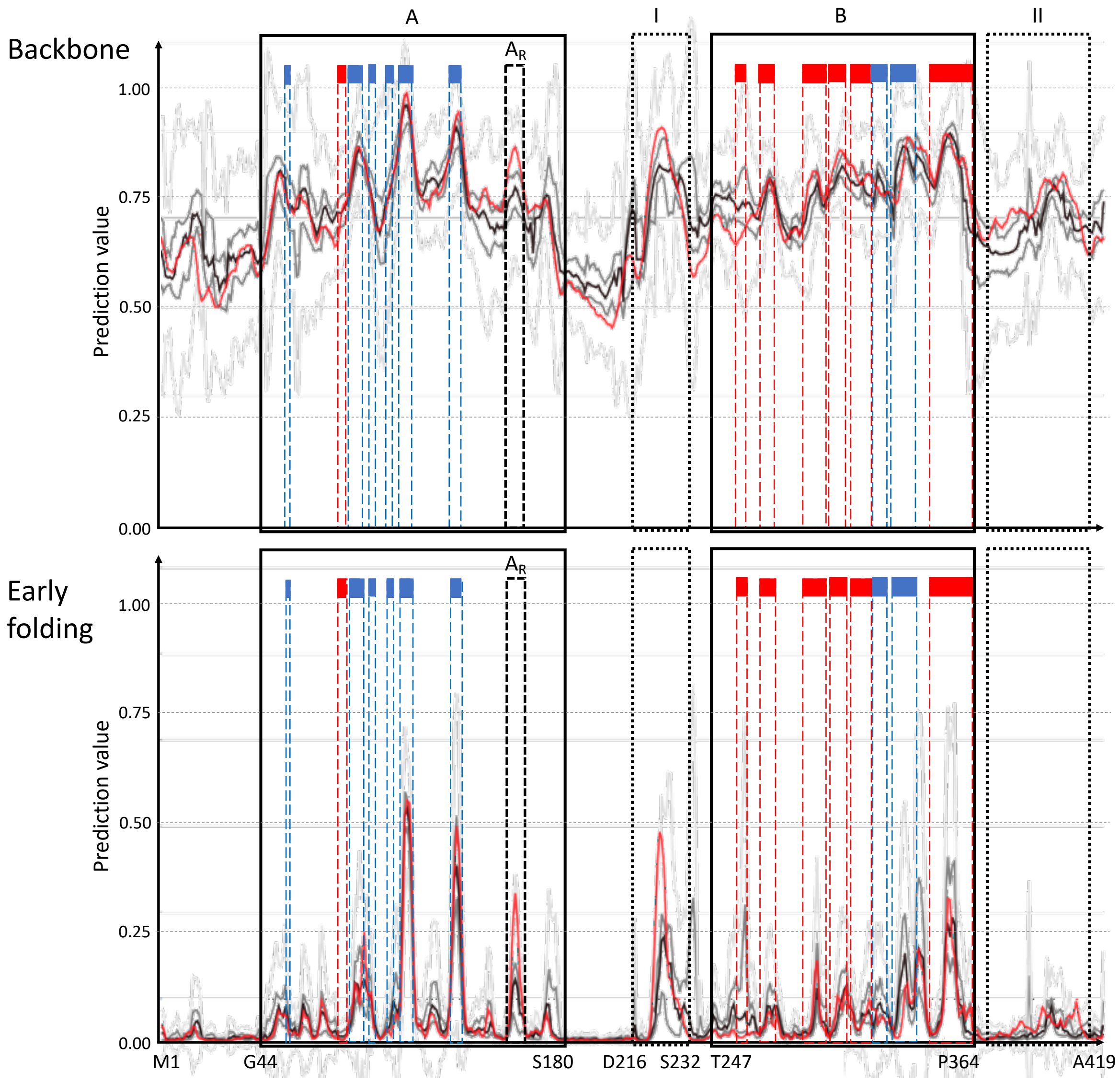


**Figure S1**: Spread of biophysical predictions based on the multiple sequence alignment (MSA) for the P0DTC9 SARS-CoV-2 protein for backbone dynamics (top) and early folding (bottom). The red line is the actual prediction for the P0DTC9 sequence, the black line represents the median value of the distribution for the MSA column corresponding to that amino acid residue of P0DTC9, the dark grey lines represent the first/third quartile, the light grey lines the outlier range. The secondary structure elements and regions of interest are indicated as in Figure 1.

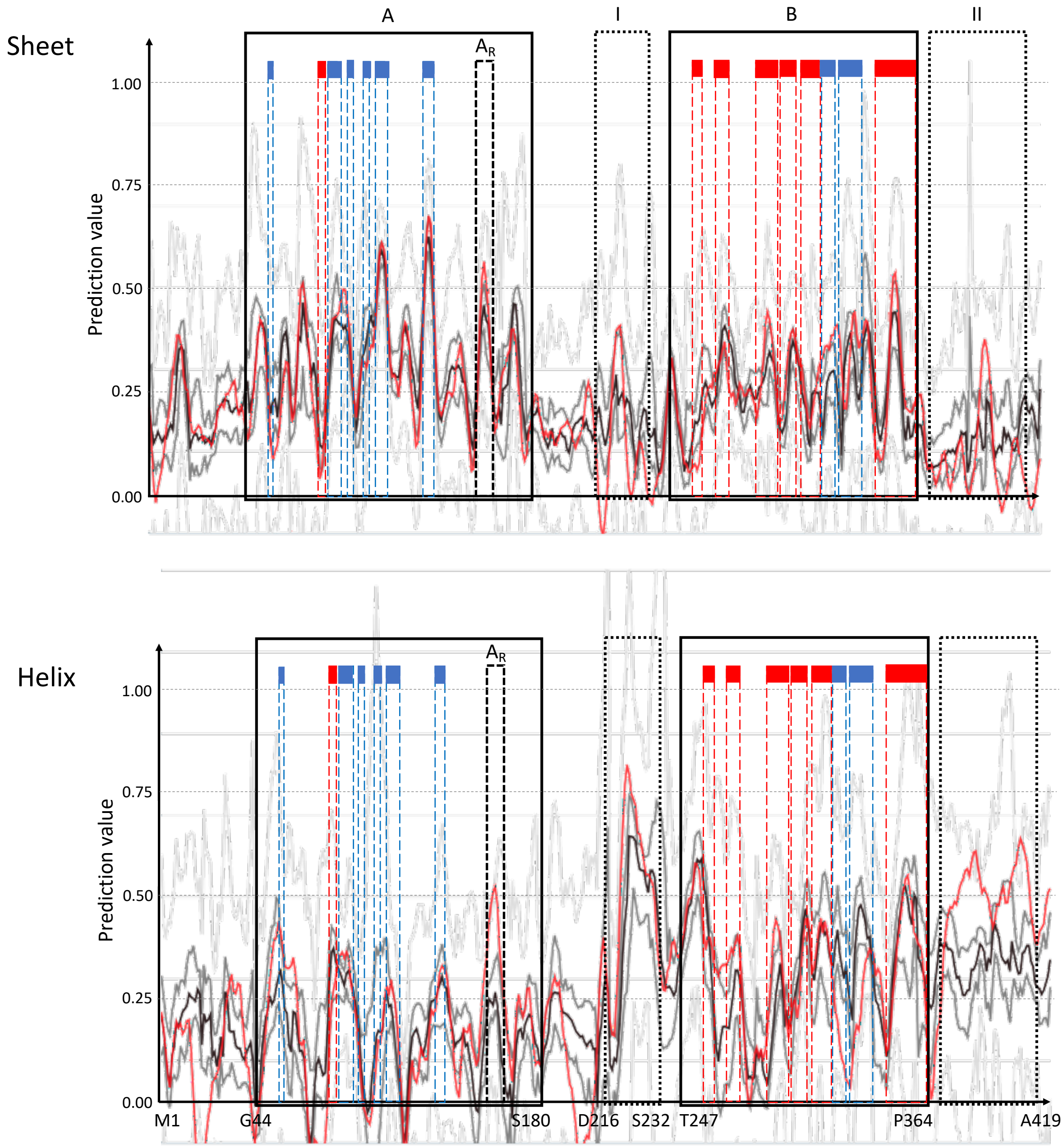


**Figure S2**: Spread of biophysical predictions based on the multiple sequence alignment (MSA) for the P0DTC9 SARS-CoV-2 protein for sheet (top) and helix (bottom) propensity. The red line is the actual prediction for the P0DTC9 sequence, the black line represents the median value of the distribution for the MSA column corresponding to that amino acid residue of P0DTC9, the dark grey lines represent the first/third quartile, the light grey lines the outlier range. The secondary structure elements and regions of interest are indicated as in Figure 1.
